## Supplemental Material for "Appearance of green tea compounds in plasma following acute green tea consumption is modulated by the gut microbiome in mice"

**Supplemental Table 1. Top amplicon sequence variants in fecal samples of low complexity microbiome mice.** Table shows the relative abundance and taxonomic assignment of the 6 highest relative abundance amplicon sequence variants in the low complexity microbiome (LCM) samples. Greengenes taxonomy was classified using a naïve Bayes classifier, whereas higher resolution taxonomic assignment was performed using basic local alignment search tool (BLAST) against the National Center for Biotechnology Information (NCBI) 16S rRNA gene database. No taxonomic assignment from Greengenes was given below the genus level, and all amplicon sequence variants shown were assigned to the phylum Firmicutes. Abbreviations: BLAST, basic local alignment search tool; NCBI, National Center for Biotechnology Information.

| Mean relative abundance in LCM samples | Top BLAST (NCBI 16S database) | % BLAST match (NCBI) | Order (Greengenes) | Family (Greengenes) | Genus (Greengenes) | Amplicon sequence variant hash |
| --- | --- | --- | --- | --- | --- | --- |
| 47.7% | *Clostridioides difficile* ATCC 9689 | 100 | Clostridiales | Peptostrepto-coccaceae | N/A | 4a4bf3c04e 4c0ccf17122 4cc4208df6e |
| 10.7% | *Zhenhengia yiwuensis* strain NSJ-12 | 100 | Clostridiales | Lachno-spiraceae | *Epulo-piscium* | 643166f2ee 8d59d8ad7c 5386f17a8293 |
| 10.2% | *Turicibacter sanguinis* strain MOL361 | 100 | Turicibacterales | Turici-bacteraceae | *Turicibacter* | a37e6c3829 6d5b9dfaffb 767794f138a |
| 7.7% | *Clostridium jeddahitimonense* strain CL-2 | 100 | Clostridiales | Clostri-diaceae | N/A | 7533d51b73 2866f837e6b 481c2cd9dba |
| 3.7% | *Pygmaiobacter massiliensis* strain Marseille-P3336 | 99.5 | Clostridiales | Rumino-coccaceae | *Anaerofilum* | 3200e29a3d 4643e9ca909 7da44f86f28 |
| 3.3% | *Niameybacter massiliensis* strain Mt14 | 99.1 | Clostridiales | Lachno-spiraceae | *Epulo-piscium* | 27c96da0e9 97cff6329b4 80cf03ad86b |

**Supplemental Table 2. Green tea compounds associated with microbiome unweighted UniFrac composition.** Table shows results of PERMANOVA tests run for each green tea compound to test their association with phylogenetic composition of the mouse fecal microbiomes. *p* values were corrected across all compounds, using the Benjamini-Hochberg false discovery rate correction. The log-fold change between the mean humanized versus low complexity microbiome (LCM) mice is also shown, where a positive value indicates higher compound abundance in the plasma of humanized mice, and a negative value indicates higher compound abundance in the plasma of LCM mice. Only compounds with an uncorrected *p* value < 0.1 are shown. Compounds ending in “.<number>” indicate that there were multiple compounds with that annotation, which were treated as separate compounds in the analysis. Compounds with the naming convention C<#> were unannotated. Abbreviations: BH, Benjamini-Hochberg false discovery rate correction; LCM, low complexity microbiome mice; HU, humanized microbiome mice; PC, Phosphatidylcholine; CL, Cardiolipin; MDGD, Monogalactosyldiacylglycerol; PI, Phosphatidylinositol; PA, Phosphatidic acid; DG, Diglyceride.

| **Compound** | **Raw *p*** | **BH *p*** | **Log_2_(HU/LCM)** |
| --- | --- | --- | --- |
| Montecristin | 0.001 | 0.292 | -1.38 |
| C1287 | 0.002 | 0.357 | 1.09 |
| Shikonofuran D | 0.004 | 0.357 | 5.36 |
| Acetylagmatine | 0.004 | 0.357 | -5.97 |
| PC(38:4) | 0.005 | 0.357 | 0.52 |
| C846 | 0.007 | 0.357 | -2.57 |
| C864 | 0.007 | 0.357 | -1.33 |
| Ethyl-p-methoxycinnamate | 0.007 | 0.357 | -1.17 |
| C943 | 0.008 | 0.357 | 0.56 |
| gamma-Glutamyl-alanine | 0.008 | 0.357 | -0.53 |
| Crotonic acid | 0.010 | 0.357 | -0.52 |
| Lactiflorin | 0.010 | 0.357 | -2.79 |
| C1000 | 0.010 | 0.357 | -2.19 |
| Mandelonitrile | 0.011 | 0.357 | 4.63 |
| C1461 | 0.012 | 0.357 | 0.76 |
| Leucocyanidin | 0.012 | 0.357 | -0.44 |
| C883 | 0.013 | 0.357 | -0.71 |
| C386 | 0.016 | 0.372 | -0.91 |
| PC(38:4) | 0.016 | 0.372 | 0.22 |
| Enanderinanin G.2 | 0.017 | 0.372 | 1.36 |
| Ixabepilone | 0.017 | 0.372 | -0.51 |
| C458 | 0.018 | 0.372 | -1.26 |
| CL(82:16) | 0.019 | 0.372 | 0.35 |
| Hericene B | 0.020 | 0.372 | -0.61 |
| N-Acetylglutamic acid | 0.020 | 0.372 | -0.39 |
| MGDG(36:6) | 0.020 | 0.372 | -0.22 |
| 22:3-Glc-Stigmasterol | 0.023 | 0.416 | 0.76 |
| Aspartic Acid | 0.028 | 0.449 | -0.67 |
| Gentian Violet | 0.028 | 0.449 | -0.68 |
| C987 | 0.028 | 0.449 | 0.96 |
| Ixabepilone.1 | 0.030 | 0.460 | -0.29 |
| cholesteryl beta-D-glucoside | 0.032 | 0.460 | -0.34 |
| C1085 | 0.032 | 0.460 | 0.45 |
| L-Pyridosine | 0.033 | 0.460 | -1.17 |
| C1436 | 0.034 | 0.462 | -0.64 |
| PI(38:6) | 0.035 | 0.462 | 0.30 |
| 2alpha,5alpha,10beta-Triacetoxy-14beta-propionyloxytaxa-4(20),11-diene | 0.036 | 0.468 | -0.03 |
| C1775 | 0.041 | 0.507 | -0.80 |
| Dibenzothiophene | 0.042 | 0.507 | -0.51 |
| Methyl guaia-1(10),11-dien-15-carboxylate | 0.043 | 0.507 | -0.92 |
| PC(36:3) | 0.043 | 0.507 | 0.26 |
| DG(22:4n6/0:0/20:4n3) | 0.045 | 0.507 | -0.69 |
| Dieporeticenin.1 | 0.047 | 0.507 | -0.94 |
| C729 | 0.047 | 0.507 | -0.66 |
| C1657 | 0.048 | 0.507 | -1.32 |
| Minwanensin | 0.049 | 0.507 | -0.74 |
| PA(34:6) | 0.054 | 0.548 | -1.04 |
| C428 | 0.055 | 0.548 | 0.57 |
| 7,8-Dihydroparasiloxanthin | 0.057 | 0.554 | -1.25 |
| Methyl beta-D-galactoside | 0.058 | 0.554 | -0.37 |
| DG(36:4) | 0.064 | 0.581 | 0.45 |
| Montecristin.1 | 0.064 | 0.581 | -0.99 |
| C756 | 0.064 | 0.581 | -0.44 |
| C1221 | 0.068 | 0.585 | -0.23 |
| C1509 | 0.070 | 0.585 | 0.21 |
| PA(35:2) | 0.070 | 0.585 | 0.39 |
| Volemitol | 0.073 | 0.585 | 1.57 |
| C1319 | 0.074 | 0.585 | 0.66 |
| C1130 | 0.074 | 0.585 | 0.43 |
| N2-Methylguanine | 0.074 | 0.585 | 1.47 |
| Rotundifoline | 0.076 | 0.585 | -0.42 |
| PC(42:6) | 0.077 | 0.585 | -0.52 |
| C1563 | 0.078 | 0.585 | 1.07 |
| C1936 | 0.078 | 0.585 | 0.34 |
| C493 | 0.081 | 0.599 | 0.58 |
| C221 | 0.086 | 0.625 | -0.35 |
| Spiramycin | 0.089 | 0.635 | -0.66 |
| PI(38:5) | 0.092 | 0.649 | 0.42 |
| DG(38:5) | 0.097 | 0.660 | 0.80 |
| C420 | 0.099 | 0.660 | -0.64 |

**Supplemental Table 3. Green tea compounds associated with microbiome phylogenetic diversity.** Table shows results of fixed effect regressions (with cluster robust standard errors, clustered by donor microbiome ID) testing the association between phylogenetic alpha diversity of the mouse fecal microbiomes and each individual green tea compound. *p* values were corrected across all compounds, using the Benjamini-Hochberg false discovery rate correction. The estimate column refers to the slope of the regression line, where green tea compounds were z-score transformed to allow comparison of effect sizes between compounds. Only compounds with an uncorrected *p* value < 0.1 are shown. Compounds ending in “.<number>” indicate that there were multiple compounds with that annotation, which were treated as separate compounds in the analysis. Compounds with the naming convention C<#> were unannotated. Abbreviations: BH, Benjamini-Hochberg false discovery rate correction; PC, Phosphatidylcholine; CL, Cardiolipin; MDGD, Monogalactosyldiacylglycerol; PI, Phosphatidylinositol; PA, Phosphatidic acid; DG, Diglyceride.

| **Compound** | **Raw *p*** | **BH *p*** | **Estimate** |
| --- | --- | --- | --- |
| CL(82:16) | <0.001 | <0.001 | 2.76 |
| Lactiflorin | <0.001 | <0.001 | -2.91 |
| C846 | <0.001 | <0.001 | -2.90 |
| Acetylagmatine | <0.001 | <0.001 | -3.67 |
| Aspartic Acid | <0.001 | 0.008 | -2.38 |
| 7,8-Dihydroparasiloxanthin | <0.001 | 0.012 | -1.52 |
| Montecristin | <0.001 | 0.014 | -1.70 |
| C1287 | 0.001 | 0.047 | 3.32 |
| C864 | 0.002 | 0.088 | -1.82 |
| C1094 | 0.002 | 0.100 | -1.94 |
| C1461 | 0.006 | 0.212 | 2.51 |
| Shikonofuran D | 0.007 | 0.219 | 2.99 |
| C883 | 0.007 | 0.219 | -1.80 |
| C458 | 0.008 | 0.219 | -2.70 |
| Ethyl-p-methoxycinnamate | 0.008 | 0.219 | -1.98 |
| C987 | 0.008 | 0.219 | 2.22 |
| gamma-Glutamyl-alanine | 0.009 | 0.220 | -2.55 |
| Ixabepilone | 0.010 | 0.236 | -1.91 |
| PI(38:6) | 0.011 | 0.236 | 2.11 |
| Mandelonitrile | 0.012 | 0.253 | 2.49 |
| MGDG(36:6) | 0.013 | 0.253 | -2.19 |
| C2106 | 0.015 | 0.263 | -0.94 |
| Leucocyanidin | 0.015 | 0.263 | -1.82 |
| Minwanensin | 0.015 | 0.263 | -1.76 |
| Ixabepilone.1 | 0.017 | 0.265 | -1.31 |
| C294 | 0.019 | 0.265 | 1.87 |
| C1436 | 0.019 | 0.265 | -2.00 |
| Crotonic acid | 0.019 | 0.265 | -2.29 |
| PC(38:4) | 0.019 | 0.265 | 2.58 |
| C943 | 0.019 | 0.265 | 2.51 |
| Oleanolic acid.2 | 0.022 | 0.284 | -0.85 |
| C1716 | 0.023 | 0.284 | 1.47 |
| Hericene B | 0.023 | 0.284 | -1.52 |
| Broussonin C | 0.023 | 0.284 | 1.35 |
| Dieporeticenin.1 | 0.025 | 0.301 | -1.76 |
| Methyl guaia-1(10),11-dien-15-carboxylate | 0.027 | 0.320 | -1.38 |
| C1353 | 0.029 | 0.332 | 1.40 |
| C1278 | 0.030 | 0.337 | 1.30 |
| N-Acetylglutamic acid | 0.032 | 0.344 | -1.38 |
| 2alpha,5alpha,10beta-Triacetoxy-14beta-propionyloxytaxa-4(20),11-diene | 0.033 | 0.344 | -2.02 |
| C1563 | 0.034 | 0.344 | 1.54 |
| DG(22:4n6/0:0/20:4n3) | 0.037 | 0.359 | -1.65 |
| C493 | 0.037 | 0.359 | 1.46 |
| PA(34:6) | 0.038 | 0.359 | -1.41 |
| PC(38:4) | 0.040 | 0.366 | 2.21 |
| Enanderinanin G.2 | 0.040 | 0.366 | 2.17 |
| 3-Teracrylmelazolide A | 0.042 | 0.371 | -1.58 |
| 2,7-Dideacetyl-2,7-dibenzoyl-taxayunnanine F | 0.043 | 0.378 | 1.21 |
| C1085 | 0.045 | 0.386 | 2.29 |
| Methyl beta-D-galactoside | 0.046 | 0.390 | -0.90 |
| Valganciclovir | 0.053 | 0.425 | 1.35 |
| C1936 | 0.054 | 0.425 | 1.88 |
| PC(36:3) | 0.055 | 0.425 | 1.95 |
| N2-Methylguanine | 0.056 | 0.425 | 1.82 |
| C2327 | 0.058 | 0.425 | 1.23 |
| PA(39:1) | 0.059 | 0.425 | 1.09 |
| C428 | 0.059 | 0.425 | 1.83 |
| L-Pyridosine | 0.059 | 0.425 | -2.29 |
| C1407 | 0.060 | 0.425 | 1.41 |
| Methyl 16-epiquillate.1 | 0.071 | 0.494 | -1.03 |
| C1772 | 0.075 | 0.514 | -0.57 |
| PI(38:4) | 0.076 | 0.515 | 1.45 |
| C1706 | 0.080 | 0.525 | -1.26 |
| C77 | 0.080 | 0.525 | -1.41 |
| C629 | 0.083 | 0.530 | 1.51 |
| Volemitol | 0.083 | 0.530 | 1.48 |
| C1518 | 0.085 | 0.530 | -1.32 |
| C239 | 0.086 | 0.531 | 1.62 |
| Rubescensin C | 0.093 | 0.537 | -1.07 |
| Oleanolic acid | 0.095 | 0.537 | -0.80 |
| C1221 | 0.096 | 0.537 | -1.75 |
| alpha-Tocopherol succinate | 0.097 | 0.537 | -0.93 |
| DG(38:5) | 0.098 | 0.537 | 1.55 |
