## Supplemental Methods for "Appearance of green tea compounds in plasma following acute green tea consumption is modulated by the gut microbiome in mice"

Preliminary Acute Feeding Study: Because limited low complexity microbiome (LCM) mice were available, a small pilot utilizing wild type (WT) mice was conducted to determine the optimal time and dose to collect samples. These WT mice were gavaged with 0, 5, 20, or 50 mg/kg green tea (GT, n=4 per group) and plasma samples were collected at 2, 4, and 48 hours post-gavage. Plasma was analyzed using mass spectrometry-based metabolomics and data analyzed to evaluate GT compounds appearing in the plasma. Methods are detailed below and in the main manuscript. Using a dose of 50 mg/kg resulted in multiple GT compounds with peak amounts at 2 hours post-gavage (data not shown). Therefore, the subsequent study used this dose and time.

Mouse cohort description: Due to the limited availability of LCM mice, the experiments were divided into three cohorts as follows: Cohort 1 comprised 3 LCM mice and 5 sets of humanized (HU) mice, Cohort 2 experiments were conducted approximately 1 month later and comprised 3 LCM mice and 3 sets of HU mice, Cohort 3 experiments were conducted approximately 1 month after Cohort 2 and comprised 3 LCM mice and 2 sets of HU mice (**Supplemental Methods Table 1**).

**Supplemental Methods Table 1: Mouse Cohorts**.

| **Mouse ID** | **Group** | **Microbiome #** | **Cohort** |
| --- | --- | --- | --- |
| GF-N June | Low complexity microbiome | None | 1 |
| GF-R June | Low complexity microbiome | None | 1 |
| GF-B June | Low complexity microbiome | None | 1 |
| GLD074-N | Humanized | 74 | 1 |
| GLD074-R | Humanized | 74 | 1 |
| GLD086-N | Humanized | 86 | 1 |
| GLD086-B | Humanized | 86 | 1 |
| GLD063-N | Humanized | 63 | 1 |
| GLD063-R | Humanized | 63 | 1 |
| GLD066-N | Humanized | 66 | 1 |
| GLD066-B | Humanized | 66 | 1 |
| GLD089-R | Humanized | 89 | 1 |
| GLD089-B | Humanized | 89 | 1 |
| GF-1 July | Low complexity microbiome | None | 2 |
| GF-2 July | Low complexity microbiome | None | 2 |
| GF-3 July | Low complexity microbiome | None | 2 |
| GLD116-N | Humanized | 116 | 2 |
| GLD116-R | Humanized | 116 | 2 |
| GLD120-N | Humanized | 120 | 2 |
| GLD120-R | Humanized | 120 | 2 |
| GLD090-N | Humanized | 90 | 2 |
| GLD090-R | Humanized | 90 | 2 |
| GF-N Aug | Low complexity microbiome | None | 3 |
| GF-R Aug | Low complexity microbiome | None | 3 |
| GF-L Aug | Low complexity microbiome | None | 3 |
| GLD101-N | Humanized | 101 | 3 |
| GLD101-R | Humanized | 101 | 3 |
| GLD113-N | Humanized | 113 | 3 |
| GLD113-R | Humanized | 113 | 3 |

*Metabolomics:* Frozen plasma samples and four GT extracts for untargeted metabolomics analysis were thawed on ice and prepared for analyses as previously described^1, 2, 3, 4^. Briefly, 50 µL of experimental samples were spiked with 5 µL each of hydrophobic and hydrophilic spike mixes, proteins were precipitated from study samples and small molecules were extracted from supernatants using an organic liquid-liquid extraction technique with methyl tert-butyl ether (MTBE) and water. The lipid fractions were reconstituted in 100 µL of methanol and the aqueous fractions were reconstituted in 50 µL 5:95 acetonitrile:water (v/v). As many samples were less than 50 µL, volumes of the spike mixes and reconstitution solvents were adjusted to maintain the same sample:spike:reconstitution volume ratio of 50:5:100 for the lipid fraction (v/v/v) and 50:5:50 for the aqueous fraction (v/v/v). For example, if only 40 µL plasma was available, 4 µL of the spike mixes were added and after liquid-liquid extraction, the samples were reconstituted in 80 µL methanol (lipid fraction) and 40 µL 5:95 acetonitrile:water (aqueous fraction).

Liquid Chromatography Mass Spectrometry-based Metabolomics: Aqueous and lipid fractions were analyzed by liquid chromatography mass spectrometry (LCMS) using published methods^2, 5, 6^. Both fractions were analyzed using an Agilent 6545 liquid chromatography-quadrupole time-of-flight mass spectrometer (LC-QTOF-MS) (Agilent Technologies, Santa Clara, CA, USA). The hydrophobic fraction of both plasma and GT samples were analyzed using reverse phase chromatography with an Agilent Zorbax Rapid Resolution HD (RRHD) SB-C18, 1.8µL (2.1 mm x 100mm) analytical column. The injection volume was 4 µL with a flow rate of 0.7 mL/min. The mobile phase A included water with 0.1% formic acid and the mobile phase B included 60:36:4 2-propanol:acetonitrile:water with 0.1% formic acid. The lipid gradient was as follows: 0-0.5 min 70% B, 0.5-7.42 min 70-100% B, 7.42-10.4 min 100% B, 10.4-10.5 min 100-70% B, 10.5-15.1 70% B. The autosampler tray temperature was set to 4 °C and the column temperature was set to 60 °C. The MS conditions were as previously described, including the following: positive ionization mode, scan rate of 2 spectra/s, mass range m/z 75–1700, drying gas temperature 300 °C and flow rate of 12.0 L/min, nebulizer pressure 35 psi, sheath gas temperature 275 °C, sheath gas flow 12 L/minute, skimmer 65 V, capillary voltage 3500 V, fragmentor 100 V, and reference masses 121.050873 and 922.009798 (Reference mix, Agilent Technologies).

The hydrophilic (aqueous) fraction was analyzed using a Zorbax SB-AQ C18, 5 µm (2.1 mm x 100 mm) analytical column. The injection volume was 5 µL with a flow rate of 0.25 mL/min. Mobile phase A included 0.1% formic acid in water with 0.1% InfinityLab Deactivator Additive, and mobile phase B included 0.1% formic acid 90% aqueous Acetonitrile with 0.1% InfinityLab Deactivator Additive. The aqueous gradient was as follows for positive mode: 0-3.0 min 1.8% B, 3-10 min 1.8-54% B, 10-15 min 54-90% B, 15-20 min 90% B, 20-20.1 min 90-1.8% B, 20.1-25 min 1.8% B. The autosampler tray temperature was set to 4 °C and the column temperature was set to 30 °C. The QTOF was run in positive ionization mode, scan rate 2.0 spectra/second, mass range m/z 50-1700, gas temperature 300 °C, gas flow 12.0 L/min, nebulizer 35 psi, skimmer 65 V, capillary voltage 3500 V, fragmentor 100 V, reference masses 121.050873 and 922.009798 (Reference mix, Agilent Technologies).

Metabolomics Quality Control: Quality control (QC) included analysis of purchased plasma (BioIVT, Westbury, NY), spiked and prepared alongside study samples. Fifteen (15) stable isotope labelled authentic standards, 9 hydrophobic (from SPLASH Lipidomix mix, Avanti Polar Lipids, Alabama, U.S.A.) and 6 hydrophilic (**Supplemental Methods Table 2**), were spiked into experimental and QC samples to monitor sample preparation and instrumental variations.

**Supplemental Methods Table 2: Aqueous and Lipid Standards.** These %CVs are typical for plasma, whereby endogenous compounds can cause interference/ion suppression and variability in standards. Abbreviations: CV, coefficient of variation.

| **Name** | **Vendor** | **%CV Plasma quality control samples** | **%CV Experimental samples** |
| --- | --- | --- | --- |
| Creatinine-d3 | CDN Isotopes | 5.96 | 23.17 |
| Adenosine (Ribose-13C5) | CIL | 7.72 | 31.77 |
| L-Acetylcarnitine-d3:HCL | CIL | 6.51 | 33.78 |
| Lactate-d3 (Sodium salt) | CIL | 4.90 | 20.58 |
| L-Carnitine-d3:HCL | CIL | 5.29 | 31.43 |
| Valine-d8 | Sigma | 4.33 | 10.86 |
| 15:0-18:1(d7) PC | Avanti | 0.86 | 12.6 |
| 15:0-18:1(d7) PE | Avanti | 3.51 | 10.9 |
| 15:0-18:1(d7) PG | Avanti | 7.14 | 12.1 |
| 18:1(d7) LPC | Avanti | 5.05 | 12.1 |
| 18:1(d7) LPE | Avanti | 6.48 | 14.4 |
| 18:1(d7) MG | Avanti | 22.54 | 14.1 |
| 15:0-18:1(d7) DG | Avanti | 12.85 | 20.4 |
| 15:0-18:1(d7)-15:0 TG | Avanti | 19.91 | 50.5 |
| 18:1(d9) SM | Avanti | 5.03 | 10.1 |

Analysis of authentic standards in BioIVT plasma QC samples was performed following each LCMS batch (i.e. daily) by generating extracted ion chromatograms (EIC) and by evaluating total ion chromatograms (TIC) using Mass Hunter Qualitative software (Agilent Technologies). For this experiment, all aqueous fractions were analyzed in a single batch in a single day. All lipid samples were also analyzed in a single batch in a single day, but on a different day than the aqueous samples. There was low/no sample injection for two lipid instrument QCs due to insufficient volume in the autosampler vial; therefore, these were removed from further QC analysis and two injections of an experimental sample were used to ensure instrument reproducibility (n=3 total injections). Standards in QC samples were evaluated visually for reproducible peak shape and height, as well as retention time (RT) by overlaying EICs. No outliers were found by this visual inspection and subsequent analysis of RT for both lipid and aqueous standards determined < 0.1 min shifts, which our laboratory has found is low and acceptable for plasma metabolomics.

Following visual and RT inspections, the signal height for each standard was analyzed in Profinder (v.10.0, Agilent Technologies) using batch targeted feature extraction (BTFE). The coefficient of variation (%CV) for each standard was calculated for QC and experimental samples for each instrument batch (QC and experimental samples were kept separate for %CV calculations). For this experiment, %CVs for 8 of the 9 lipid standards was <20% for QC samples, <10% for 3 repeated injections of a single sample at batch beginning and end, and <20% for experimental samples. These results are typical for this method and sample type.

In addition, for aqueous fractions, %CVs for the 6 aqueous standards was <8% for QC samples and <35% for experimental samples.  Some variation is to be expected in experimental samples due to experimental conditions and biological differences. However, %CVs over 20% for experimental samples is high based on our experience. BioIVT plasma QC samples were also compared to experimental samples using principal components analysis (PCA) to determine if outliers or batch effects were present.  A PCA performed for this purpose using raw aqueous data (2,679 compounds), prior to any filters or normalization, shows separation of QC and experimental plasma as expected. Two clusters of experimental samples are also seen in aqueous data, which did not appear to be due to preparation day or cohort, as evidenced by samples from each cohort and prep day in each cluster (Supplemental Figure 1, left panel). As demonstrated by the instrument QC results, no batch effects were due to LCMS. Because LCM and HU samples from each cohort were present in each cluster and no technical issues were found, aqueous data was extracted as described and evaluated as described in the main text, with a batch effect taken into account. Note that clustering of raw lipid data (3,435 compounds) using PCA (Supplemental Figure 1, right panel) showed expected distribution of QC and experimental samples.

Effects of cohort on composition of the fecal microbiome, plasma GT metabolome, and plasma GT lipidome: Mouse cohort was not significantly correlated with differences in microbiome composition (PERMANOVA on centroids *R*^2^ = 0.14, pseudo-*F* = 0.8, *p* = 0.70), and was thus not adjusted for in most statistical analyses. Additionally, our PERMANOVA results for microbiome, GT lipids, and GT aqueous compounds were consistent between stratified (only permuted within each mouse cohort) and unstratified (permutated across all mice) analyses.

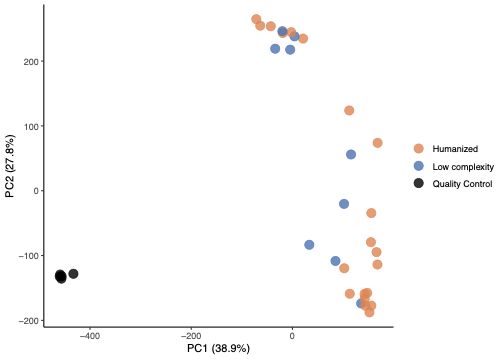

**Supplemental Figure 1**: Using whole metabolome data (rather than green tea compounds only), PCA of experimental and quality control plasma samples showed 3 clusters for aqueous samples (A) and the expected 2 clusters for lipid samples (B). Quality control samples cluster away from study samples, as expected. The 2 clusters of aqueous plasma compounds were due to different sample preparation days. Color distinguishes samples from low complexity microbiome mice, humanized microbiome mice, and quality control samples.

Preliminary experiment using mice not fed GT: A preliminary experiment measuring the plasma metabolomes from LCM mice and HU mice not fed green tea was conducted to determine baseline differences. As with the main study, mice were placed on an irradiated, low polyphenol diet (TD.97184, Envigo, Indianapolis, IN) at the time of weaning (3 weeks of age) through 6 weeks of age. At 4 weeks of age, mice were orally gavaged with 200 µL of either human fecal slurry (100 mg stool homogenized in 1 mL reduced PBS in an anaerobic chamber) or PBS for LCM controls. For this experiment, a total of 16 mice were used; 6 mice were gavaged with vehicle and 10 mice were gavaged with the same 10 distinct human fecal slurries as described in the main text (i.e. 1 microbiome per mouse). At two weeks post-gavage, plasma was collected as described in the main text. Metabolomics of plasma was performed as described in the main text and above using LC/MS. Database searching was performed in ID Browser (Agilent Technologies) as described in the main text. Briefly, match criteria included mass (<10 ppm window), isotope peak abundance and spacing, and a match score cutoff of 60. Compounds were first searched against in-house standards databases, containing over 600 compounds. For in-house standards databases retention time was required for a match with a tolerance window of 0.3 minutes. Unannotated compounds were then searched against FooDB, METLIN, Human Metabolome Database (HMDB), Kyoto Encyclopedia of Genes and Genomes (KEGG), and Lipid Maps. For compounds matched to more than one database, the hit with the highest score was assigned as the annotation. Compounds that matched to a database entry represent a Metabolomics Standards Initiative level 3 identification, based on the proposed minimum reporting by Sumner, et al. and hence compound names are considered tentative.

Matching compounds from the GT mice and the non-GT controls: For each compound found in the plasma of mice fed GT (n=624 compounds), the monoisotopic ion was used to mine the plasma metabolomics data from the GF and HU mice not fed GT. Briefly, extracted ion chromatograms are manually generated for each green tea compound using non-GT plasma LC/MS data employing a targeted data extraction strategy using Qualitative Analysis (Agilent). For each GT compound extracted ion chromatogram, the MS spectrum was manually inspected to verify the targeted ion was present as a monoisotopic peak.  Only quantifier/qualifier peaks detected within +/- 0.2 minutes of the retention time recorded in samples are retained. Briefly, if the monoisotopic ion for a GT compound was present in the non-GT data (as a monoisotopic peak only) and the retention time was within 0.2 min, it was considered a match. In all cases the mass error was less than 5 ppm.
